# Biosynthetic Glycan Labeling

**DOI:** 10.1101/2021.07.01.450741

**Authors:** Victoria M. Marando, Daria E. Kim, Phillip J. Calabretta, Matthew B. Kraft, Bryan D. Bryson, Laura L. Kiessling

**Affiliations:** Department of Chemistry, Massachusetts Institute of Technology, Cambridge, Massachusetts 02139, United States; Department of Chemistry, University of Wisconsin Madison, Madison, Wisconsin 53706, United States; Department of Biological Engineering, Massachusetts Institute of Technology, Cambridge, Massachusetts 02139, USA; Ragon Institute of MGH, MIT, and Harvard, Cambridge, Massachusetts 02139, USA

## Abstract

Glycans are ubiquitous and play important biological roles, yet chemical methods for probing their structure and function within cells remain limited. Strategies for studying other biomacromolecules, such as proteins, often exploit chemoselective reactions for covalent modification, capture, or imaging. Unlike amino acids that constitute proteins, glycan building blocks lack distinguishing reactivity because they are composed primarily of polyol isomers. Moreover, encoding glycan variants through genetic manipulation is complex. Therefore, we formulated a new, generalizable strategy for chemoselective glycan modification that directly takes advantage of cellular glycosyltransferases. Many of these enzymes are selective for the products they generate yet promiscuous in their donor preferences. Thus, we designed reagents with bioorthogonal handles that function as glycosyltransferase substrate surrogates. We validated the feasibility of this approach by synthesizing and testing probes of D-arabinofuranose (D-Ara*f*), a monosaccharide found in bacteria and an essential component of the cell wall that protects mycobacteria, including *Mycobacterium tuberculosis*. The result is the first probe capable of selectively labeling arabinofuranose-containing glycans. Our studies serve as a platform for developing new chemoselective labeling agents for other privileged monosaccharides. This probe revealed an asymmetric distribution of D-Ara*f* residues during mycobacterial cell growth and could be used to detect mycobacteria in THP1-derived macrophages.

---

Monomer-selective bioconjugation reactions have transformed the way in which we study biomolecules, affording molecular-level insight into structure, function, localization and dynamics.^1-3^ Proteins exhibit a large degree of functional group variation with respect to their constituent amino acids. This functional group diversity has been the driver of methodology development in the protein bioconjugation field. In contrast, glycans and their component sugars cannot easily be distinguished from one another on the basis of complementary reactivity, as the structural diversity of glycans derives predominantly from the stereo- and constitutional isomerism of polyol monomers (**Figure 1A**). Thus far, the systematic interrogation of the structure-function relationships that underpin molecular signaling at the cell surface has been limited by our inability to chemically perturb glycans with the requisite degree of precision and selectivity.

**Figure 1.**
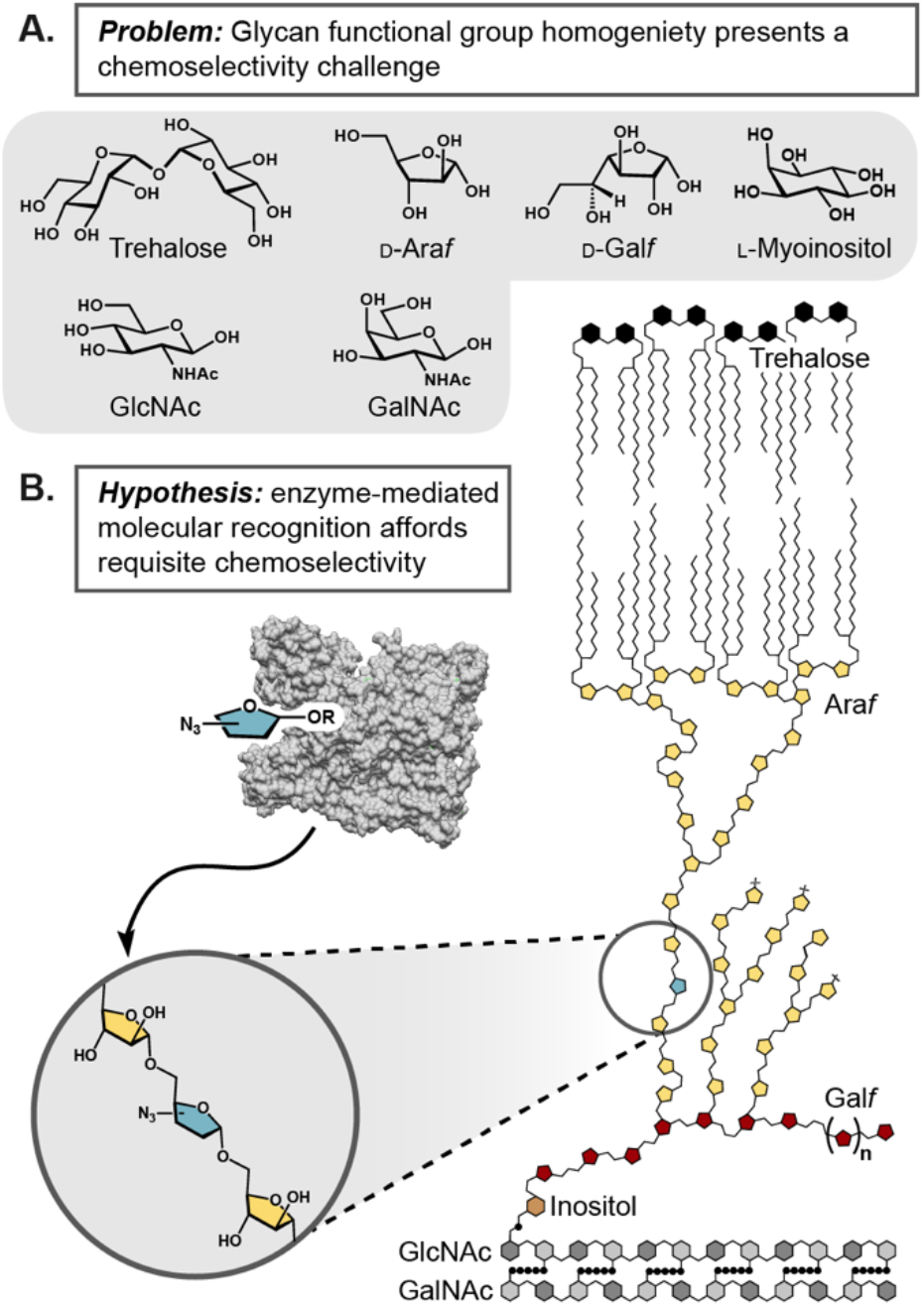
(A) Monosaccharides, the building blocks of glycans, derive their identity on the basis of polyol stereo- and constitutional isomerism. (B) The core cell wall structure of Mycobacteria and Corynebacteria is comprised of six unique monomers. This structure, termed the mycolyl-arabinogalactan-peptidoglycan complex (mAGP) is a dense glycolipid matrix that protects cells from environmental stressors and antibiotics. Biosynthetic incorporation leverages the evolved affinity of extracellular glycosyltransferases for specific monosaccharides to introduce chemoselective modifications into complex cell surface glycans.

In the absence of a viable chemical approach to site-selective glycan labeling, metabolic engineering has been used to modify and study cell surface glycans in eukaryotic systems.^4-7^ This technique traditionally relies on the cellular uptake of non-natural monosaccharides, followed by extensive biosynthetic processing to nucleotide-sugar analogs. As nucleotide-sugars can serve as donors for cytosolic glycosyltransferases, these intermediates are incorporated into growing glycan chains that are subsequently exported to the cell surface. This strategy is effective in eukaryotes; however, the adaptation of metabolic incorporation to prokaryotes has been historically challenging.^8-13^ Mammals utilize 35 unique monosaccharide building blocks, while bacteria employ over 600.^14-15^ The structural diversity of bacterial monosaccharides and glycans necessitates a complex, and often poorly understood carbohydrate metabolism.^16-17^ As a result, small molecule probes that are reliant on metabolic machinery for extensive biosynthetic processing and display are unlikely to be incorporated in a predictable or site-selective manner.

In formulating a chemical strategy to address the problem of selective glycan modification, we began from the assumption that within a complex glycan, unique sugar monomers can be best distinguished from one another on the basis of molecular recognition, not complementary reactivity.^18^ Fortunately, catalysts with the requisite selectivity already exist in nature -the endogenous glycosyltransferases. These biocatalysts have an evolved selectivity for a specific small molecule substrate donor and a specific polysaccharide acceptor (**Figure 1B**).^19-23^

In this report, we describe the development and application of a novel approach to chemoselective glycan bioconjugation. This strategy, termed biosynthetic incorporation, leverages the activity of endogenous extracellular enzymes using substrate surrogates (**Figure 2**). This biocatalytic manifold can side-step challenges associated with small molecule-based glycan bioconjugation and metabolic engineering by exploiting the intrinsic selectivity of the target biocatalysts.

**Figure 2.**
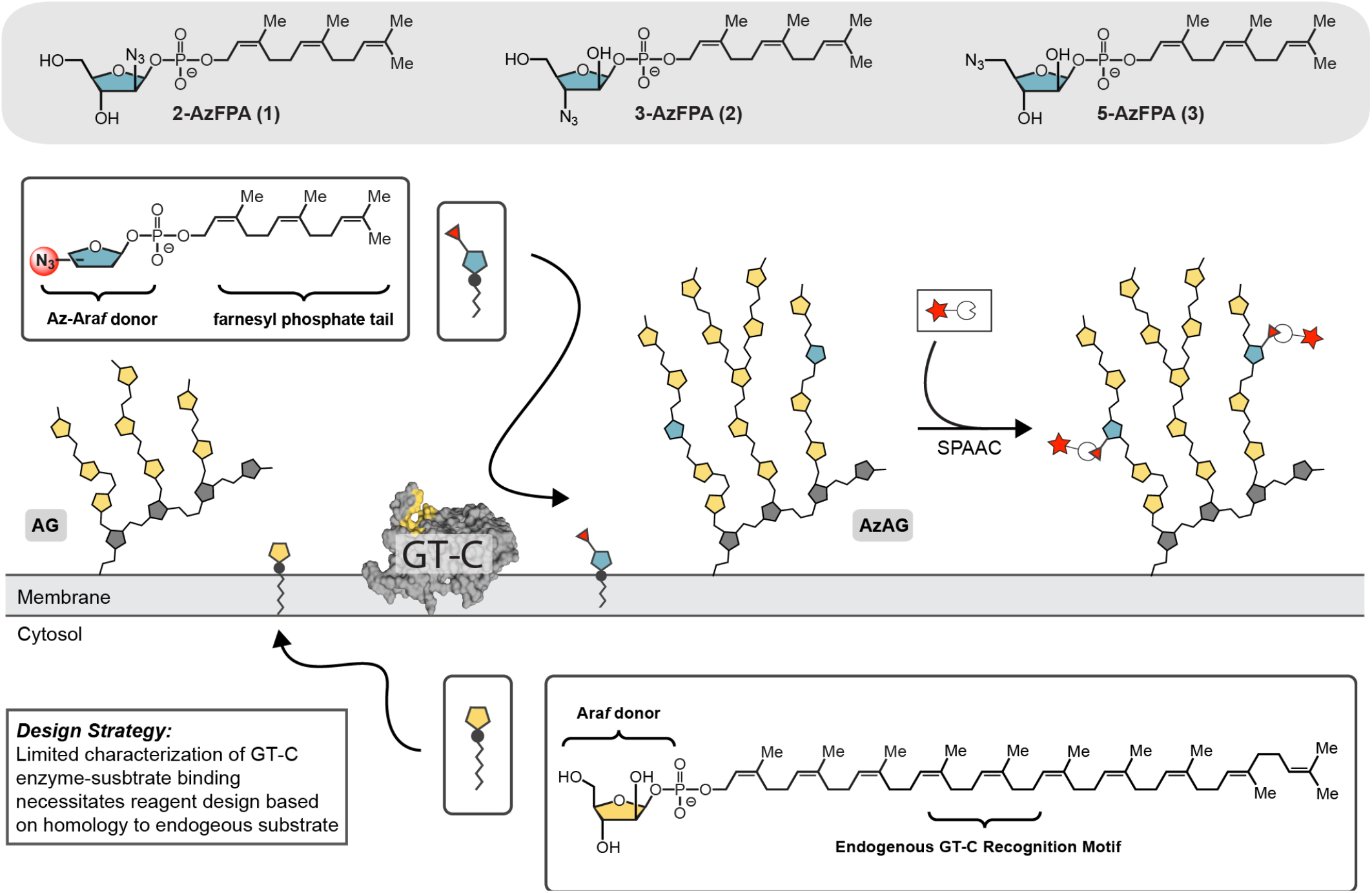
To harness the catalytic activity of arabinofuranosyltransferase GT-Cs for bioconjugation, an azide-modified substrate surrogate was designed based on structural homology to the endogenous D-Ara*f* donor (DPA). Three azide regioisomers were produced (**1-3**). Exogenous delivery of AzFPA was expected to result in substrate incorporation, which could subsequently be detected and quantified using SPAAC-mediated fluorophore conjugation.

To demonstrate the feasibility of our chemoselective labeling strategy, we targeted D-arabinofuranose (D-Ara*f*). This arabinose isomer is not found in humans but is present in microbes. D-Ara*f* is an essential component of the cell wall of the order Mycobacteriales.^24-25^ Although many constituents of this order are benign, *Mycobacterium tuberculosis* (*Mtb*), *Corynebacterium diphtheriae*, and *Mycobacterium leprae* are notorious human pathogens.^26-28^ These organisms utilize D-Ara*f* for the construction of the core glycolipid component of their cell wall, the mycolyl-arabinogalactan-peptidoglycan complex (mAGP).^29^ Within the mAGP, arabinan is thought to play an important role in maintaining the overall structural integrity of the cell envelope. Indeed, its biosynthesis is the target of the front-line antituberculosis drug ethambutol.^30^ D-Ara*f* residues are well suited to demonstrate the capabilities of our strategy. First, no methods to selectively visualize D-Ara*f* are known. Second, the biosynthesis of the activated sugar proceeds through late-stage epimerization of the C2 hydroxyl from the ribose-phospholipid to the corresponding arabinose-phospholipid donor rather than other sources of free arabinose.^31^ Consequently, metabolic engineering approaches are unlikely to result in specific D-Ara*f* labeling.^32^

In mycobacteria, D-Ara*f* is incorporated into cell surface glycans through the action of integral membrane glycosyl-transferases (GT-Cs).^19-23^ In contrast to the nucleotide-sugar substrates of cytosolic glycosyltransferases, these GT-Cs are transmembrane enzymes that recognize polyprenyl phosphate-linked sugar donors.^33^ Our group previously demonstrated that the non-natural glycolipid donor (*Z,Z*)-farnesyl phosphoryl-β-D-arabinofuranose (FPA) is a surrogate for the extended (C_55_) endogenous arabinofuranose donor decaprenyl phosphoryl-β-D-arabinofuranose (DPA) in *C. glutamicum* and *M. smegmatis*.^34^ This lipid substitution facilitates practical exogenous reagent delivery, as extended polyprenyl glycosides are poorly soluble and form micelles.^22, 35^ Here, we employ this non-natural (*Z,Z*)-farnesyl phosphoryl donor scaffold as a vehicle to introduce azide-modified D-Ara*f* derivatives into the mAGP. Although the endogenous monosaccharide donor and polysaccharide acceptors for cell wall arabinosylation were well characterized, few 3D structures for the target arabinosyltransferase enzymes had been reported.^36^ We reasoned that the efficiency of incorporation could vary between different isomers. Accordingly, we focused on preparing and evaluating all three possible regioisomeric azido-(*Z,Z*)-farnesyl phosphoryl-β-D-arabinofuranose (AzFPA) derivatives.

The AzFPA regioismers were synthesized from commercially available arabinofuranose and ribofuranose monomers. The key azide functionality was installed through nucleophilic displacement or nucleophilic epoxide opening.^37^ We appended the (*Z,Z*)-farnesyl recognition motif and then removed the protecting groups to afford the target compounds (See Supporting Information). The synthetic routes were optimized to provide access to the desired substrate surrogates for microbiological studies.

The collection of AzFPA isomers was evaluated for incorporation in *C. glutamicum* and *M. smegmatis* using fluorescent labeling and flow cytometry (Figure 3). Bacteria were cultured to mid-logarithmic phase with each AzFPA isomer, washed to remove unassociated probe, and treated with AF647-conjugated dibenzocyclooctyne (DBCO) to install the fluorophore via a strain-promoted azide-alkyne click reaction (SPAAC).^38^ At the indicated dose, no change in bacterial viability was observed due to probe treatment (See Supporting Information).

**Figure 3.**
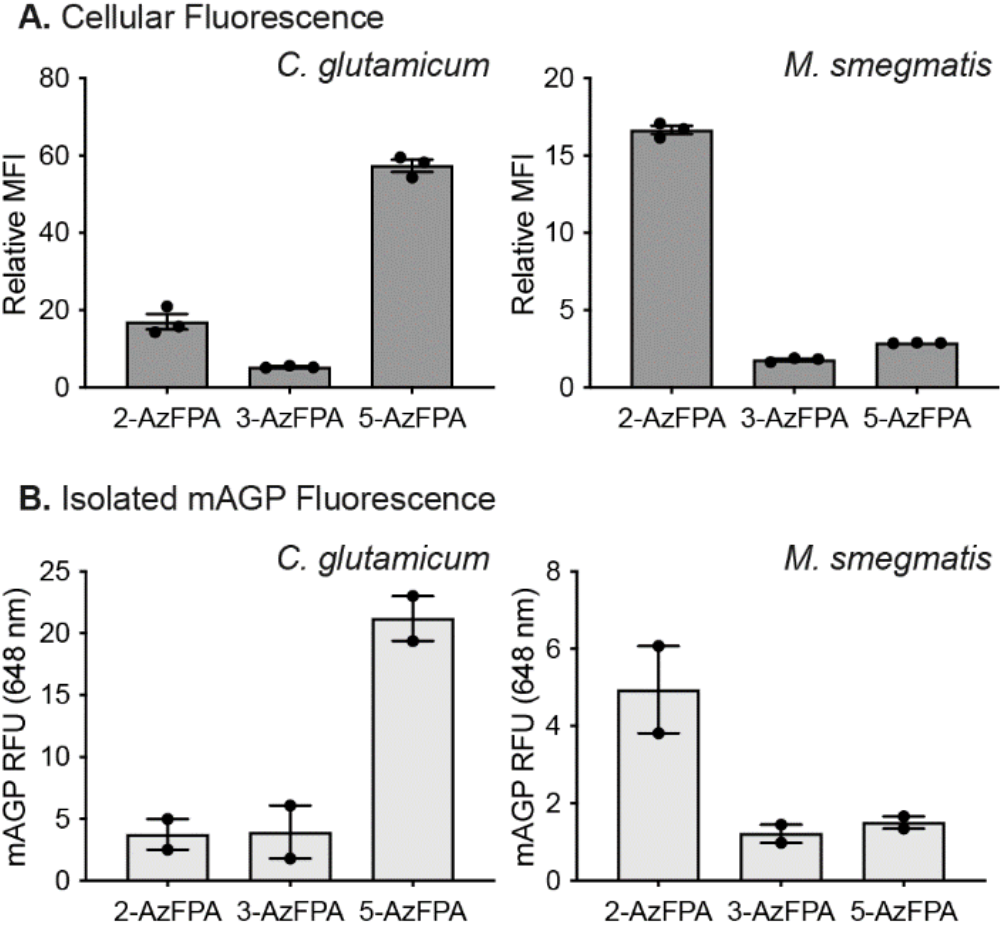
(A) Flow cytometry of AzFPA (250 uM) labeled *C. glutamicum* and *M. smegmatis* treated with DBCO-AF647. MFI was calculated using the geometric mean and plotted relative to a dye only control. Error bars denote the standard error of the mean of three replicate experiments. (B) Fluorescence emission (633 nm) from isolated mAGP from AzFPA (250 uM) labeled *C. glutamicum* and *M. smegmatis* reacted with DBCO-AF647. Error bars denote the standard error of the mean of two replicate experiments.

Analysis of fixed samples by flow cytometry revealed increased fluorescence of cells treated with 2-AzFPA (**1**) and 5-AzFPA (**3**). The 2-azido isomer exhibited the brightest staining in *M. smegmatis* while the 5-azido derivative led to more *C. glutamicum* labeling (**Figure 3A**). The 3-AzFPA (**2**) derivative afforded minimal staining. The observed fluorescence due to labeling compound **1** and **3** was not the result of non-specific staining, as *E. coli*, an organism lacking arab-inose-containing cell surface glycans, exhibited no staining (**Figure S1**). The observed selectivity is consistent with recently disclosed structural data of the *M. smegmatis* arabinofuranosyltransferase EmbA bound to the endogenous arabinose donor DPA, determined by cryoelectron microscopy (Cryo-EM).^36^ This structure indicates there are key hydrogen-bonding contacts between the C-3 hydroxyl group of DPA within the catalytic pocket. Thus, the poor incorporation of the 3-AzFPA (**2**) regioisomer could be the result of disrupted enzyme-substrate complementarity. A finding from the experiments above is that different substrates are preferred by each species. This observation highlights the value of testing different isomers. Isomer preferences may be further tuned to enable the selective functionalization of distinct glycans within specific organisms.

To determine whether AzFPA was incorporated into the target arabinose-containing biopolymer, the mAGP was isolated. We applied a standard procedure to cells that had been modified with AzFPA and stained with AF647.^34, 39^ Fluorescence of each isolated glycopolymer fraction was used to determine the degree of incorporation (**Figure 3B**). The trends in fluorescence for mAGP modification were consistent with relative cellular fluorescence observed by flow cytometry. The reduced fluorescence intensity across samples could be attributed to the rigorous isolation protocol.

We next tested our probes in confocal fluorescence microscopy to visualize the localization of the mAGP within live cells. As before, the bacteria were cultured in the presence of AzFPA and stained with AF647-DBCO. The relative incorporation trends first detected by flow cytometry were mirrored by the intensities of fluorescence observed by microscopy; the 5-and 2-AzFPA probes afforded the most pronounced signal in *C. glutamicum* and *M. smegmatis* respectively (See Supporting Information). Taken together, these data indicate that the AzFPA probes are competent substrates for relevant glycosyltransferases and can be selectively incorporated into cell surface glycans.

Having validated the efficacy of our platform for fluorescence-based applications, we proceeded to use 2-AzFPA (**1**) to visualize cell wall biosynthesis. Live cell confocal imaging of *M. smegmatis* revealed brighter staining at the poles and septum in dividing cells (**Figure 4A**). The spatial localization of the probe was quantified across cell length. The resultant fluorescence intensity plot was consistent with previous literature reports of other cell wall probes in *M. smegmatis*, namely, 7-hydroxycoumarin-3-carboxylic acid-3-amino-D-alanine (HADA), a fluorescent D-alanine analog that is incorporated into nascent peptidoglycan and a mycolic acid probe, quencher-trehalose-fluorophore (QTF).^40-41^

**Figure 4.**
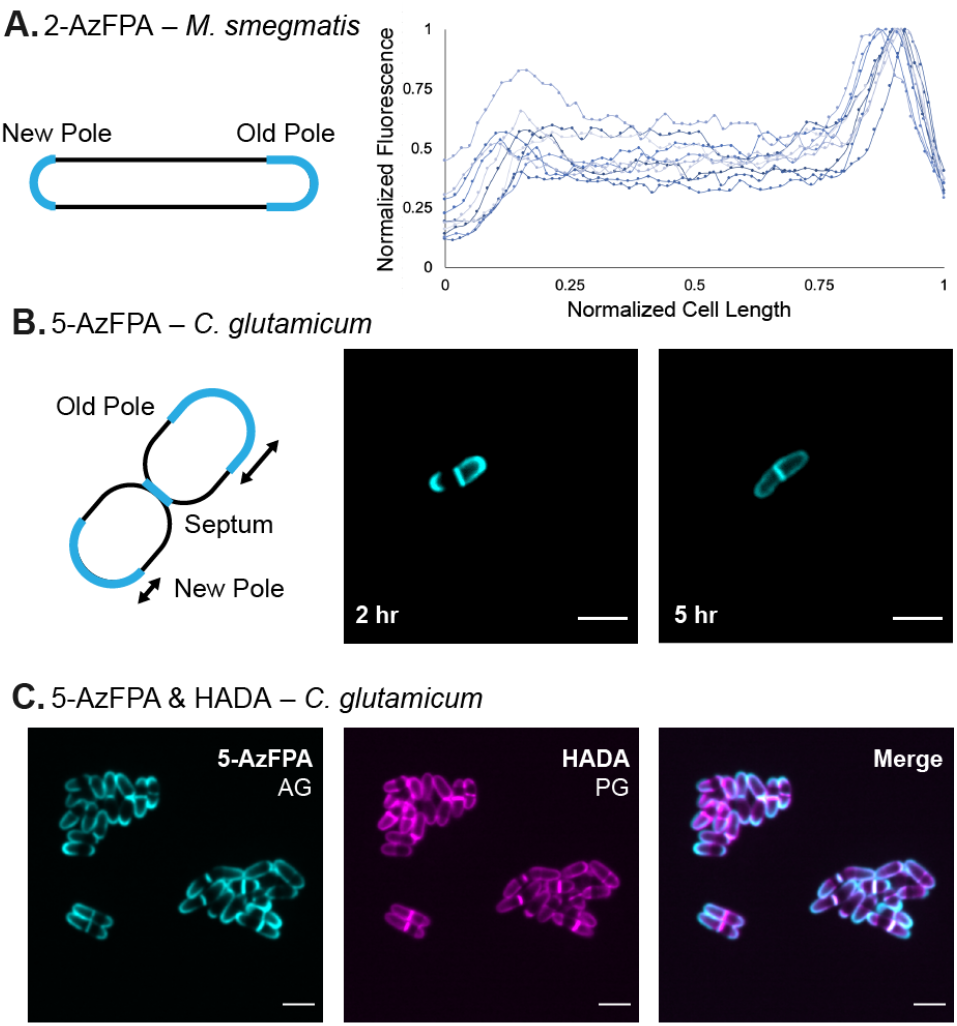
(A) Localization analysis of *M. smegmatis* grown with 2-AzFPA (250 uM). (B) Confocal fluorescence microscopy images of C. glutamicum grown with 5-AzFPA (250 uM) for 2 or 5 hours. (C) Confocal fluorescence microscopy images of C. glutamicum grown with 5-AzFPA (250 uM) and HADA (500 uM) for 2 hours. (Scale bars: 3 um).

To better understand arabinogalactan biosynthesis, we visualized 5-AzFPA (**3**) incorporation in *C. glutamicum* (**Figure 4B**). As the incorporation of our probes is contingent upon enzymatic activity, we expected to observe brighter staining in areas where cell wall biosynthesis and remodeling is most active. Like *M. smegmatis, C. glutamicum* grows asymmetrically, with growth occurring most rapidly at the old pole and slower at the new pole and septal plane.^42^ Exposure of cells to 5-AzFPA (**3**) over two doubling times afforded pronounced polar and septal staining. As cells underwent multiple division cycles after five doubling times, the staining pattern became distributed across the cell envelope. The ability of cells to continue dividing in the presence of the probes and the morphology of the bacteria indicate that no major deleterious changes to the cell envelope occur.

The asymmetry in the fluorescence pattern was similar to that observed previously for the peptidoglycan.^41^ Because the peptidoglycan serves as the base cell wall structure to which the arabinogalactan is conjugated, we anticipated that arabinogalactan and peptidoglycan assembly would coincide. To test this hypothesis, we incubated *C. glutamicum* with 5-AzFPA (**3**) and HADA to visualize the mAGP and peptidoglycan simultaneously.^41^ The fluorescent signals co-localized (**Figure 4C**). These data support our hypothesis and indicate that our probe can be used to in concert with established tools to explore cell envelope assembly.

To assess the utility of our probes in a more complex environment, we examined their utility for visualizing bacteria in a phagocytic cell. *Mtb* infection initiates from aerosol particles that enter the lungs of a host.^43^ The bacteria then recruit macrophages to the lung that phagocytose the invading pathogen.^44^ To examine whether labeled bacteria could be detected, we pre-stained *M. smegmatis* with 2-AzFPA then exposed them to THP1-derived macrophages (**Figure 5A**). Bacteria were taken up by the phagocytic cells, and the fluorescent signal was stable (**Figure 5B**). These data indicate that our probes can be used to visualize D-Ara*f* residues in more complex systems, such as infection models.

**Figure 5.**
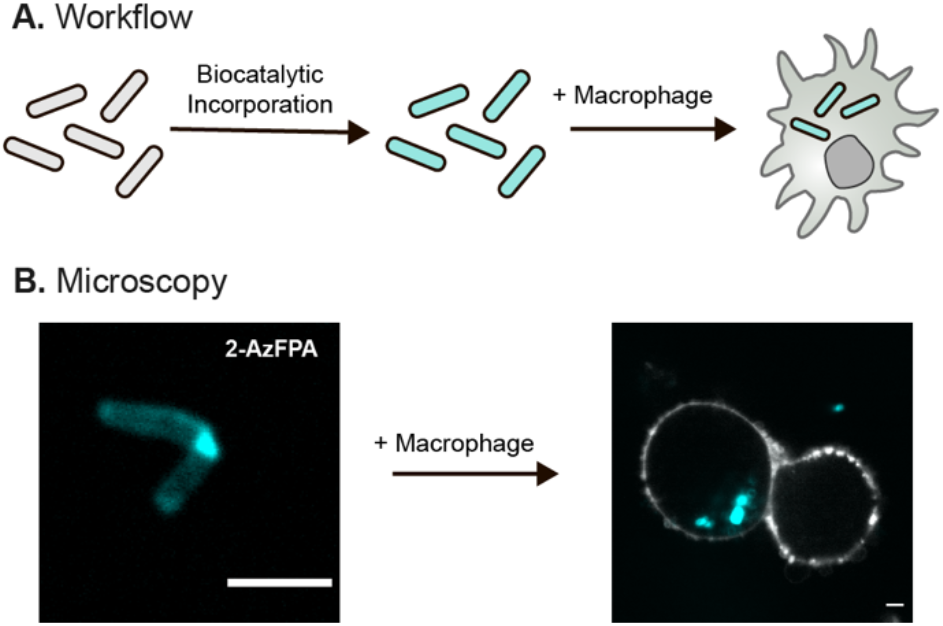
(A) Schematic of the macrophage infection work flow. First *M. smegmatis* cells are labeled with AzFPA, then infection into human derived macrophage is observed. (B) Confocal fluorescence microscopy images if *M. smegmatis* with THP1-derived macrophages. (Scale bars: 3 um).

Our findings reveal that synthetic glycolipid donors can be used for chemoselective chemical modification of cell surface glycans. Using a suite of tools for *in cellulo* D-Ara*f* functionalization, we identified probes that could give rise to species-selective glycan modification. These findings indicate that biosynthetic incorporation can be exploited to selectively modify the glycans of different species—even when these glycans are constructed from identical building blocks. The disclosed AzFPA reagents will enable new studies including the facile purification of glycans, the visualization of polysaccharide trafficking, biosynthesis, and remodeling, as well as the identification of new protein-carbohydrate interactions at the cell interface. We anticipate that the work described here will serve as a foundation for the further expansion of the biosynthetic incorporation platform to other monosaccharide components of complex glycans.

## AUTHOR INFORMATION

### Notes

The authors declare no competing financial interest.

## ACKNOWLEDGEMENT

The authors thank H. L. Hodges, R. L. McPherson and K. I. Taylor for helpful scientific discussions as well as R. L. McPherson and S. D. Brucks for their assistance in reviewing the manuscript. The authors thank the NIH-NIAID (Al-126592 to L.L.K.), the NIH Common Fund (UO1GM125288 to L.L.K.), the NIH (R01A1022553 and R01AR073252 to B.D.B.) NIH-NIGMS (F32 GM142288-01 to D.E.K.), and the NSERC (PGSD Fellow-ship for V.M.M.) for financial support.

