## Supplemental Information for "Biosynthetic Glycan Labeling"


#### Table of Contents


Supplemental Procedures....S2

Supplemental Figures....S4

Chemical Synthesis and Characterization Data....S7

Supplemental References....S20

#### Supplemental Procedures

##### Strains and Growth Conditions

For bacterial assays, the strains employed include *Mycobacterium smegmatis mc2155*, *Corynebacterium glutamicum ATCC13032* and *Escherichia coli BL21*. *M. smegmatis* was grown in Middlebrook 7H9 broth (HiMedia, Mumbai, India) supplemented with 0.2% (w/v) dextrose, 0.2% (v/v) glycerol, 0.5% bovine serum albumin (United States Biological, Salem, MA), catalase (4 mg/liter) (Sigma-Aldrich), 15 mM sodium chloride and 0.05 % (v/v) Tween 80 in a shaking incubator at 37°C. *C. glutamicum* was cultured in brain heart infusion (BHI) medium (BD, Franklin Lake, NJ) supplemented with 9% (w/v) sorbitol and *E. coli* was cultured in LB liquid medium (Sigma-Aldrich) in a shaking incubator at 30°C and 37°C, respectively. Generally, starter cultures were incubated at the desired temperature with shaking until saturation. Cells were then diluted into fresh media and grown to mid-logarithmic phase (determined by OD600 measurement on a BioMate 3S Spectrophotometer).

##### Growth Inhibition Experiments

Experiments were performed following reported procedures.[1] In brief, a saturated culture was diluted down to the desired starting OD and plated in triplicate in a Corning black 96-well plate. Probes were added at the indicated concentration from DMSO stock solutions. *M. smegmatis* were grown with shaking at 37°C for 24 h. *C. glutamicum* were grown with shaking at 30 °C for 16 h. Alamar Blue reagent (6 µL, Invitrogen) was added to each well and the plates were incubated again for 1 h at 37 °C or 30°C. The fluorescence per well was then measured on a Tecan Infinite M1000 Pro microplate reader. Monitoring of resorufin fluorescence was achieved by exciting at 570 nm ± 5 nm and detecting at 585 nm ± 5 nm. Z-position was set to 2 mm, and fluorimeter gain was optimized and then kept constant between plates. Data are reported in relative fluorescence units (RFU) normalized to untreated controls (**Figure S2**).

##### Flow cytometry and Fluorescence Microscopy

Cells were plated from saturated starter cultures in a Corning black 96-well plate inoculated from the saturated starter cultures ($OD_{600}$ = 0.05). *M. smegmatis* was cultured in Middlebrook 7H9 containing 0.2% (v/v) glycerol and 0.05 % Tween 80. *C. glutamicum* was cultured in BHIS medium containing 0.05 % Tween 80. *E. coli* was cultured in LB liquid medium containing 0.05 % Tween 80. AzFPA derivatives were added to the desired concentrations from 85 mM stocks in DMSO. Cultures were grown to mid-log ($OD_{600}$ 1.0-1.2) at 37 °C or 30 °C with shaking. Samples were immediately prepared for flow cytometry or microscopy.

Cells were pelleted for 5 min at 3000 x g. The pellets were washed with ice-cold PBS supplemented with 0.05% Tween 80 (100 µL) once. Cells were washed an additional time with PBS supplemented with 0.05% Tween 80 and 0.5% (w/v) BSA once then taken up in fresh 7H9 media supplemented with 0.5% Tween 80 (150 µL). AFDye™ 647 DBCO (Click Chemistry Tools #1302) was added from a 10 mM stock solution in DMSO to a final concentration of 500 µM. The samples were stained for 2 h rotating at 37 °C. When 1 hr and 45 mins minutes had passed, 0.1 µL (1500X dilution) of SytoBC™ Green Fluorescent Nucleic Acid Stain (ThermoFischer #S34855) was added for a 15-minute incubation period. The stained cells were pelleted for 5 min at 3000 x g. The supernatant was removed, and the pellet was washed with PBS supplemented with 0.05% Tween 80 and 0.5% (w/v) BSA twice.

For flow cytometry, stained cell pellets were taken up in 4% paraformaldehyde in PBS to be fixed at room temperature for 20 min. Following fixation, cells were pelleted then taken up in sterile PBS supplemented with 0.05% Tween 80 in flow tubes and analyzed using an Attune NxT Flow Cytometer (405 nm, 488 nm, 561 nm, and 640 nm lasers). 10,000 cells were counted at the low flow rate. Flow cytometry analysis was performed in triplicate, representative scatter plots are shown (**Figure S3**). The unstained controls were analyzed first to set gates. Data were analyzed in FlowJo (FlowJo LLC). Mean fluorescence intensity was calculated using a geometric mean.

For analysis by microscopy, stained cell pellets were taken up in 7H9 supplemented with 0.5% Tween 80 (100 uL). Each sample was spotted onto a glass-bottomed microwell dish (MatTek corporation # P35G-1.5-14-C) and covered with a pre-cooled and 0.6% (w/v) agarose pad. Images were collected RPI spinning-disk confocal microscope (100x oil immersion lens, 1.4 NA). Brightness and contrast were identically adjusted with the open-source Fiji distribution of ImageJ. Images were then converted to an RGB format to preserve normalization and then assembled into panels (**Figure S4**).

**mAGP Isolation**

Cell envelope material was extracted similar to previous protocols.[1-2] *M. smegmatis* cultures (1 mL) were inoculated from a saturated starter culture ($OD_{600}$ = 0.05) in Middlebrook 7H9 containing 0.05 % Tween 80. AzFPA derivatives were added from DMSO stock solutions to the desired concentrations (250 μM), and the cultures were grown to saturation at 37 °C with shaking. *C. glutamicum* was cultured in BHIS medium (1 mL) in a shaking incubator at 30 °C. Cells were pelleted for 5 min at 3000 x g, normalizing across samples by OD. Cells were washed and reacted with AFDye™ 647 DBCO (Click Chemistry Tools #1302) as described above. After staining, cell pellets were resuspended in lysis buffer (2% Triton X-100 in PBS) and disrupted by sonication (6 x 20 s separated by 2 min off intervals on ice). The cell lysate was then pelleted by centrifugation at 15 000 g for 15 min. The supernatant was removed and the pellet was taken up in 2% SDS in PBS and heated to 95 °C for 1 hr before pelleting as above and discarding the supernatant. The pellet was then washed with water, 80% acetone/water and then acetone. After isolation, the mAGP complex was suspended in 2% SDS in PBS and the fluorescence was detected on a Tecan M1000 plate reader. Plates were shaken for 3 s (6 mm, orbital) immediately before the well fluorescence was read ($\lambda ex$ = 648 nm ±5, $\lambda em$ = 671 ± 5 nm).

**Uptake into THP-1 cells**

The monocyte cell line THP-1 (ATCC TIB-202) was cultured in ATCC RPMI-1640 supplemented with 10% FBS, 0.05 mM 2-mercaptoethanol and P/S in a 5% $CO_2$ humidified atmosphere. To perform infection experiments, monocyte cells were differentiated into macrophages by induction with phorbol-12-myristate-13-acetated (PMA, Sigma-Aldrich). Cells were seeded in six-well cell non-treated cell culture plates in complete RPMI with 10 uM PMA. After 48 h of PMA stimulation, cells were adherent to plate and were washed with PBS and allowed to recover in complete RPMI media for another 24 h before using THP-1-derived macrophages for infection experiment. *M. smegmatis* cells were added at the indicated multiplicity of infection (MOI) and the co-cultures was incubated at 37 °C with 5% $CO_2$ for 1 h. Macrophages were then washed PBS with 1% (w/v) BSA three times and spotted onto a glass-bottomed microwell dish. Images were collected RPI spinning-disk confocal microscope (100x oil immersion lens, 1.4 NA). Brightness and contrast were identically adjusted with the open-source Fiji distribution of ImageJ. Images were then converted to an RGB format to preserve normalization and then assembled into panels.

#### Supplemental Figures

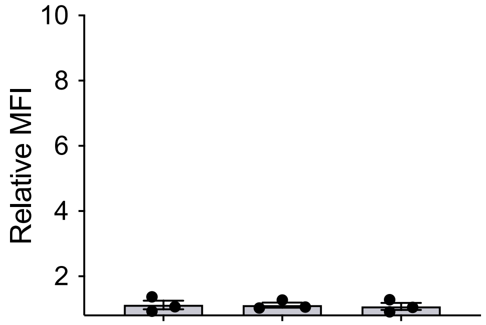


**Figure S1. Cellular Labeling in *E. coli*.** Flow cytometry of *E. coli* cultured in the presence of AzFPA (250 µM) following treatment with DBCO-AF647. MFI was calculated using the geometric mean and plotted relative to a dye only control. Error bars denote the standard error of the mean of three replicate experiments.

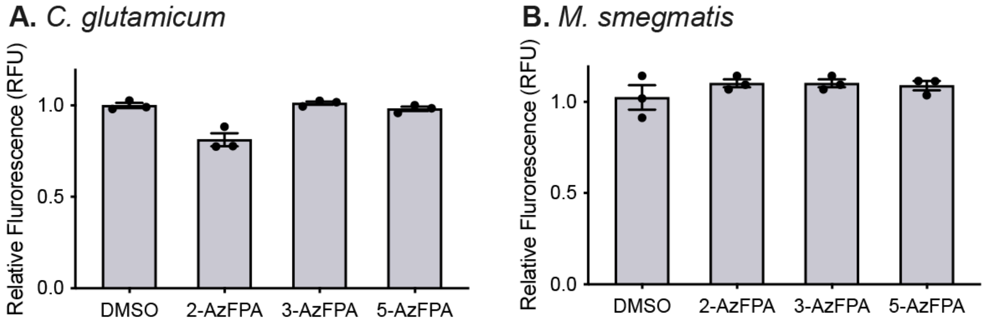


**Figure S2. Probe Viability Determined by Alamar Blue Assay.** Growth of *C. glutamicum* (A) and *M. smegmatis* (B) measured via the Alamar Blue assay in the presence of 250 µM AzFPA. The Y axis depicts the relative fluorescence compared to an untreated control sample. Error bars denote the standard error of the mean of three replicate experiments.

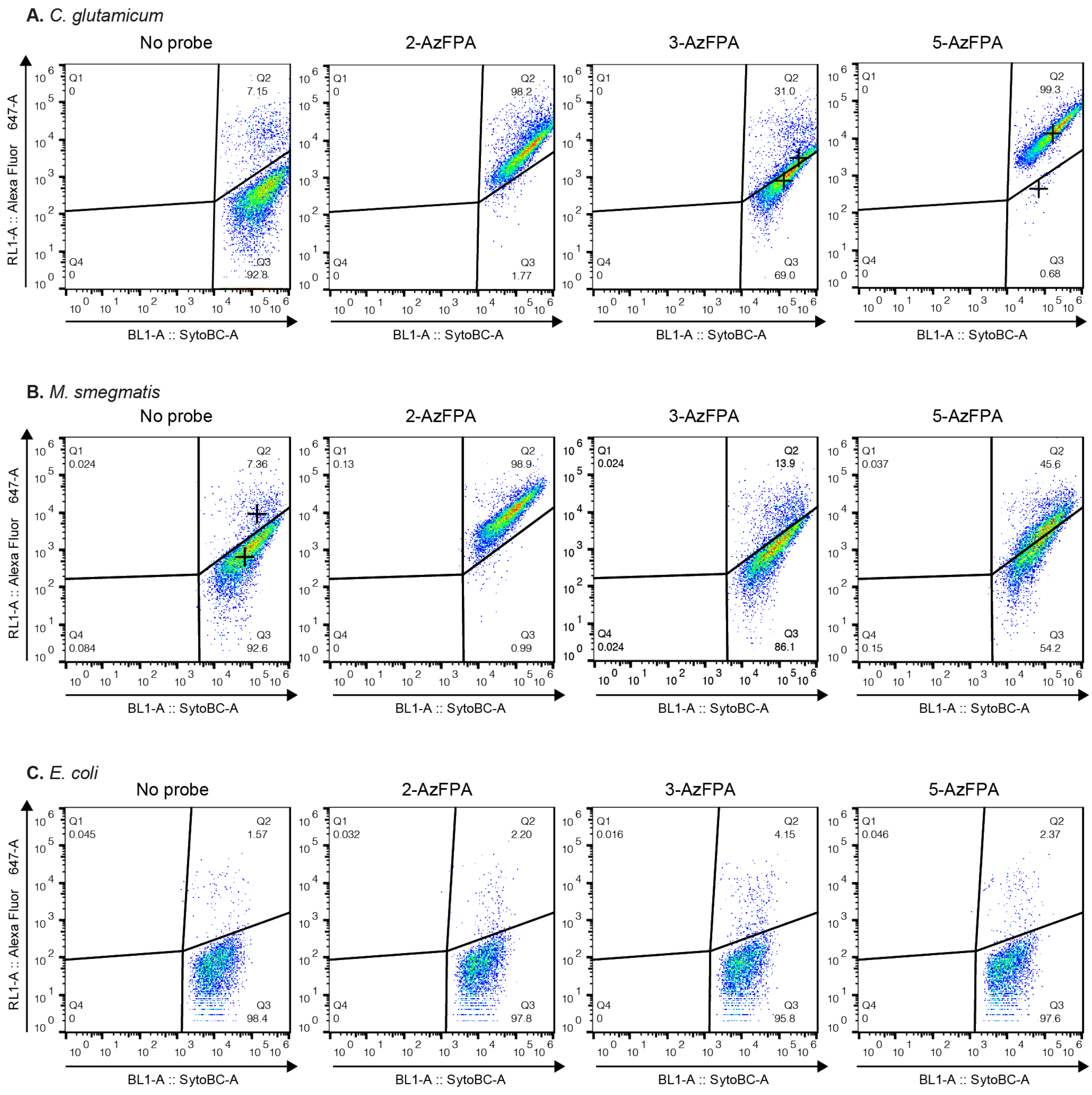


**Figure S3. Flow Cytometry Scatter Plots.** Representative scatter plots for *C. glutamicum* (A), *M. smegmatis* (B) or *E. coli* (C) treated with DBCO-AF647 in the absence or presence of AzFPA probe. Plots are representative of two independent experiments with three replicates for each strain and condition.

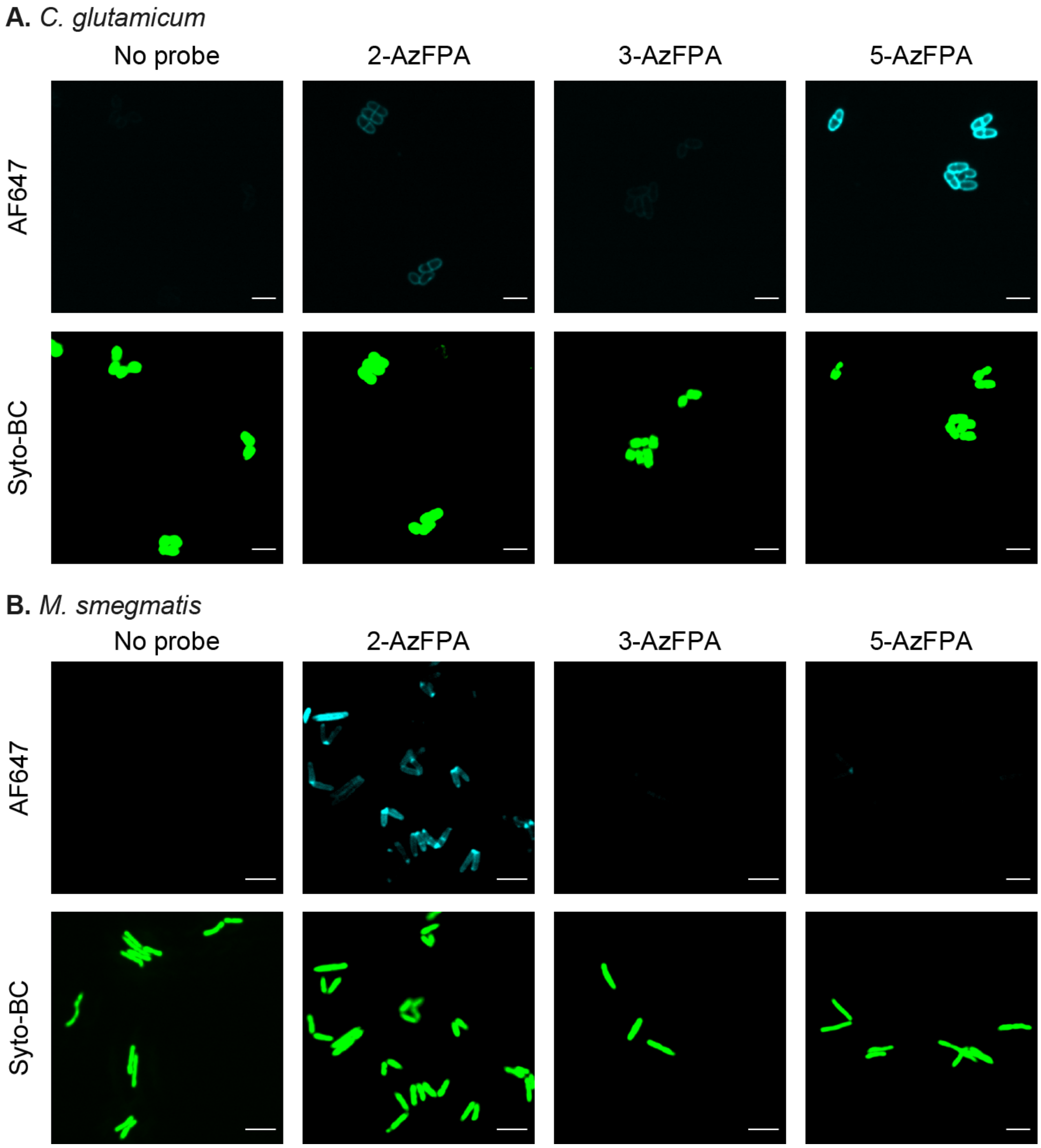


**Figure S4. Confocal Microscopy.** Fluorescence confocal microscopy of live AzFPA-labeled *C. glutamicum* (A) and *M. smegmatis* (B) reacted with DBCO-AF647. Cells were identified using nuclear stain Syto-BC. (Scale bars: 3 um).

#### Chemical Synthesis and Characterization Data

##### General information

All chemicals were purchased from Sigma Aldrich unless otherwise stated. Dry solvents were obtained from a solvent purification system (Pure Process Technologies) under argon unless otherwise stated. DMF, MTBE and pyridine were used from sure seal bottles (Sigma Aldrich) without further purification. Triethylamine was distilled from $CaH_2$ just prior to use.

Analytical thin layer chromatography (TLC) was performed on EMD Millipore TM TLC silica gel 60 F254 (glass-backed). Plates were visualized under UV light and by staining with *p*-anisaldehyde stain with charring. Flash chromatography was performed on SiliCycle ® SiliaFlash ® P60 silica gel and Biotage ® Selekt using Biotage Sfär silica cartridges.

Nuclear magnetic resonance spectra were recorded on a 300 MHz spectrometer (acquired at 300 MHz for $^{1}$H and 75 MHz for $^{13}$C), 400 MHz spectrometer (acquired at 400 MHz for $^{1}$H and 100 MHz for $^{13}$C), a 500 MHz spectrometer (acquired at 500 MHz for $^{1}$H and 125 MHz for $^{13}$C) or a 600 MHz spectrometer (acquired at 600 MHz for $^{1}$H and 151 MHz for $^{13}$C). Chemical shifts are reported relative to residual solvent peaks in parts per million ($CHCl_3$: $^{1}$H, 7.26, $^{13}$C, 77.16; MeOH: $^{1}$H, 3.31, $^{13}$C, 49.00; $C_6D_6$: $^{1}$H, 7.16, $^{13}$C, 128.06). High-resolution mass spectra (HRMS) were obtained on an electrospray ionization-time of flight (ESI-TOF) mass spectrometer. All IR spectra were taken on an FT-IR Bruker Alpha II.

### Scheme S1. Synthesis of AzFPA regioisomers

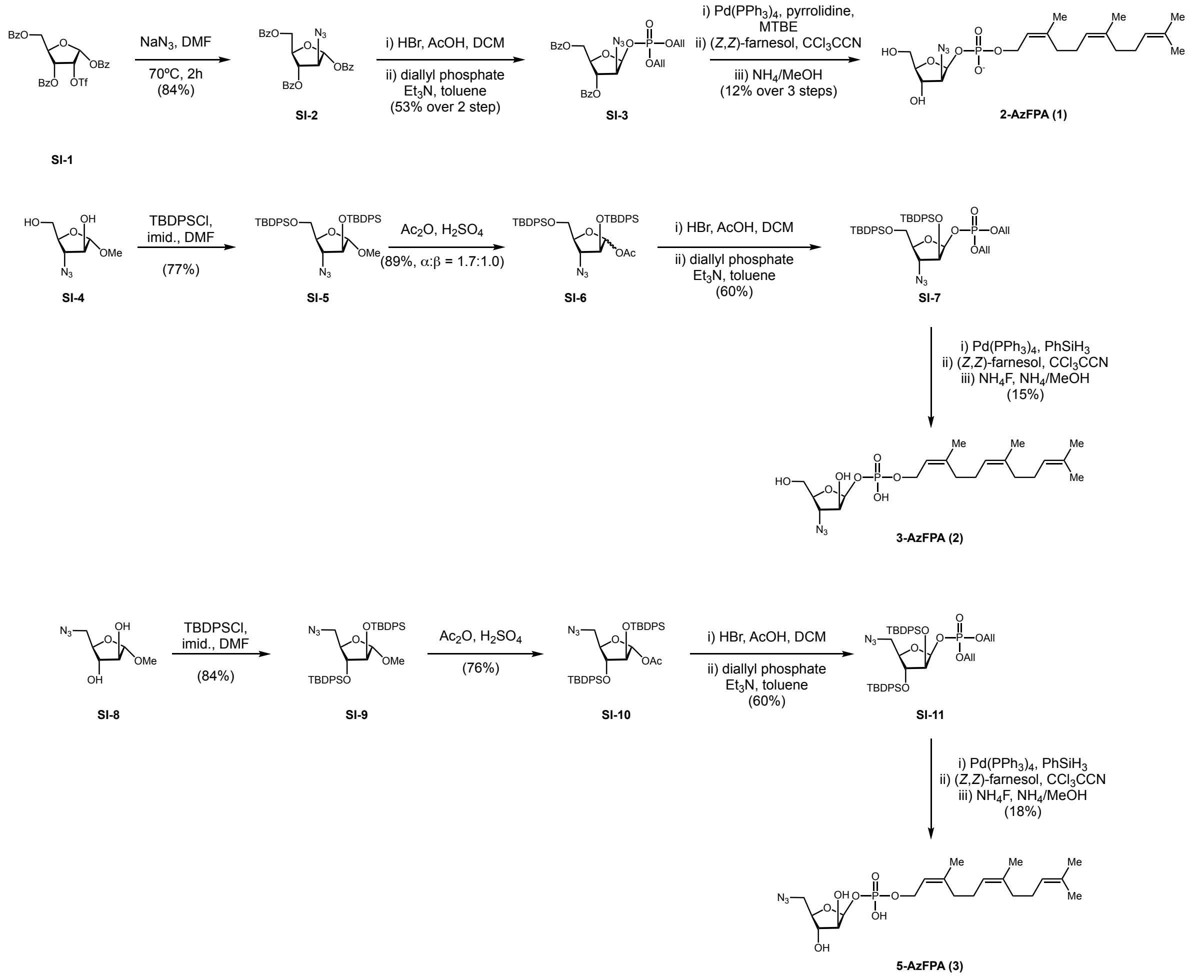

**2-azido-1,3,5-tri-*O*-benzoyl-2-deoxy-α-D-arabinofuranoside (SI-2)**.

A flame-dried, 50 mL round-bottomed flask equipped with a magnetic stir bar and a reflux condenser was charged with **SI-1** (5.00 g, 8.41 mmol, 1.0 equiv), sealed with a rubber septum, evacuated and backfilled with nitrogen three times, and placed under an argon atmosphere. **SI-1** was dissolved in anhydrous DMF (28 mL, 0.3 M) and $NaN_3$ (820.0 mg, 12.61 mmol, 1.5 equiv) was added to the resulting solution as a solid in a single portion. The reaction apparatus was transferred to an oil bath and heated to 70 °C.

After stirring at 70 °C for 5 hours, the reaction apparatus was removed from the oil bath and allowed to cool to room temperature. The reaction was quenched by pouring into sat. aq. $NaHCO_3$ (300 mL). The layers were separated and the aqueous layer was extracted with $Et_2O$ (3 × 300 mL). The combined organic layers were washed with brine (500 mL), dried over anhydrous $Na_2SO_4$, filtered, and concentrated under reduced pressure by rotary evaporation to provide a crude orange solid. Purification by flash column chromatography on silica gel (EtOAc/hexanes = 0/10 to 1/3) afforded **SI-2** (3.38 g, 84%) as a white solid.

**1H NMR (300 MHz, CDCl3)** δ 8.19 – 8.09 (m, 3H), 8.05 (dddd, J = 7.4, 3.3, 1.6, 1.0 Hz, 5H), 7.67 – 7.52 (m, 4H), 7.51 – 7.36 (m, 8H), 6.54 (s, 1H), 5.45 (ddd, J = 3.7, 1.2, 0.7 Hz, 1H), 4.84 (dt, J = 5.2, 3.9 Hz, 1H), 4.73 (dd, J = 12.0, 4.1 Hz, 1H), 4.66 (dd, J = 12.0, 5.2 Hz, 1H), 4.54 – 4.50 (m, 1H).

**13C NMR (75 MHz, CDCl3)** δ 166.03, 165.72, 164.93, 134.30, 134.16, 133.81, 130.35, 130.26, 129.81, 129.40, 129.02, 128.91, 128.89, 128.78, 128.68, 128.64, 93.35, 82.28, 79.67, 70.18, 63.67.

**IR** ($cm^{-1}$): 3062, 2956, 2101, 1717, 1597, 1577, 1491, 1451, 1312, 1242, 1103, 1086, 1067, 1020, 938

**2-azido-3,5-di-*O*-benzoyl-1-*O*-diallylphosphoryl-2-deoxy-β-D-arabinofuranoside (SI-3).**

A flame-dried, 10 mL round-bottomed flask equipped with a magnetic stir bar was charged with **SI-2** (1.00 g, 2.05 mmol, 1.0 equiv), sealed with a rubber septum, evacuated and backfilled with nitrogen three times, and placed under an argon atmosphere. **SI-2** was dissolved in anhydrous $CH_2Cl_2$ (4.1 mL, 0.5 M) and the resulting solution was cooled to 0 °C by transferring the reaction apparatus to an ice-water bath. After stirring at this temperature for 5 min, 33% HBr in AcOH (740 µl, 4.10 mmol, 2.0 equiv) was added to the reaction flask in a dropwise fashion. Following this addition, the reaction apparatus was immediately removed from the ice-water bath and allowed to equilibrate to room temperature.

After stirring for 1 hour, the stir bar was removed from the reaction flask and the reaction mixture was concentrated *in vacuo*. The resulting orange gum was dried on hi-vac for 1 hour. During this interim period, a second 25 mL round-bottomed flask equipped with a magnetic stir bar and 4 Å molecular sieves (1.00 g) was sealed with a rubber septum and evacuated. The flask was flame dried 3 times to activate the molecular sieves. Upon cooling, the reaction apparatus was evacuated and backfilled with nitrogen three times, and placed under an argon atmosphere. Diallylphosphate (730 mg, 4.10 mmol, 2.0 equiv) was transferred to the reaction flask in a single portion and dissolved in anhydrous toluene (4.1 ml, 0.5 M). Triethylamine (860 µl, 6.15 mmol, 3 equiv) was added to the reaction mixture in a dropwise fashion over the course of 5 minutes, after which the reaction was cooled to -10 °C by transferring the reaction apparatus to a NaCl-ice bath. After stirring at this temperature for 5 minutes, the glycosyl bromide intermediate was dissolved in anhydrous toluene (4.1 ml, 0.5 M). The resulting solution was transferred to the reaction flask in a dropwise fashion. Following this final addition, the reaction flask was allowed to slowly warm to room temperature.

After stirring for 10 hours, the reaction was quenched by pouring into sat. aq. $NH_4Cl$ (150 mL). The layers were separated and the aqueous layer was extracted with $Et_2O$ (3 × 1500 mL). The combined organic layers were washed with brine (300 mL), dried over anhydrous $Na_2SO_4$, filtered, and concentrated under reduced pressure by rotary evaporation to provide a crude oil. Purification by flash column chromatography on silica gel (EtOAc/hexanes = 0/10 to 1/2) afforded **SI-3** (591 mg, 53% over 2 steps) as a colorless oil.

**$^1$H NMR (300 MHz, CDCl3)** δ 8.10 – 7.98 (m, 4H), 7.61 (ddt, *J* = 8.1, 6.9, 1.4 Hz, 1H), 7.57 – 7.42 (m, 3H), 7.42 – 7.32 (m, 2H), 6.05 (dd, *J* = 5.5, 4.4 Hz, 1H), 6.02 – 5.84 (m, 1H), 5.84 – 5.77 (m, 2H), 5.38 (dq, *J* = 17.1, 1.5 Hz, 1H), 5.30 (dq, *J* = 5.8, 1.4 Hz, 1H), 5.25 (dd, *J* = 2.0, 1.0 Hz, 1H), 5.16 (dq, *J* = 10.4, 1.2 Hz, 1H), 4.79 – 4.68 (m, 1H), 4.65 – 4.49 (m, 6H), 4.22 (ddd, *J* = 8.2, 4.4, 2.3 Hz, 1H).

**13C NMR (75 MHz, CDCl3)** δ 166.22, 165.84, 134.11, 133.41, 132.37, 132.29, 132.19, 130.12, 130.00, 129.72, 128.85, 128.75, 128.58, 118.80, 118.65, 99.88, 99.81, 80.77, 77.44, 74.81, 68.82, 68.75, 68.61, 68.54, 66.26, 66.16, 65.13.

**IR** ($cm^{-1}$): 3069, 2979, 2949, 2883, 2104, 1720, 1597, 1581, 1448, 1259, 1176, 1097, 1017

**HRMS (ESI):** calc. for $C25H27N3O9P^+$ (M + H) 544.1479, found 544.1480.

**1-*O*-(*Z,Z*)-farnesylphosphoryl-2-azido-2-deoxy-β-D-arabinofuranoside, 2-AzFPA (1).**

A flame-dried, 5 mL microwave vial equipped with a magnetic stir bar and was charged with **SI-3** (300 mg, 0.55 mmol, 1.0 equiv), sealed with a rubber septum, evacuated and backfilled with nitrogen three times, and placed under an argon atmosphere. **SI-3** was dissolved in anhydrous MTBE (1.4 mL, 0.4 M) and $Pd(PPh_3)_4$ (6 mg, 1 mol%) was added to the reaction vial in a single portion. Following this addition, the reaction vial was transferred to an ice-water bath to cool to 0 °C. After 5 min, pyrrolidine (180 µl, 2.2 mmol, 4.0 equiv) was added to the reaction vial neat, in a dropwise fashion over the course of 2 minutes. After this addition, the reaction vial was removed from the ice-water bath and allowed to equilibrate to room temperature. Over the course of the reaction, the formation of a white precipitate was observed due to the insolubility of the de-allylated phosphate intermediate in MTBE.

After stirring for 2 hours, stirring was ceased and the precipitate was allowed to settle for 5 minutes. After this time, the supernatant was decanted and the remaining white precipitate was washed with MTBE (3 × 2 mL). The reaction vial was fitted with an aluminum–PTFE crimp cap and dried on hi-vac for 1 hour to remove trace MTBE and pyrrolidine. Following this, the reaction vial was evacuated and backfilled with nitrogen three times, and placed under an argon atmosphere. The crude arabinofuranosyl phosphate was dissolved in anhydrous pyridine (1.1 mol, 0.5 M). (*Z,Z*)-farnesol (367 mg, 1.65 mmol, 3.0 equiv) and trichloroacetonitrile (553 µl, 5.51 mmol, 10.0 equiv) were added to the reaction vial neat, in single portions. The resulting solution was heated to 40 °C by transferring the reaction vial to an oil bath.

After stirring at this temperature for 12 hours, the reaction vial was removed from the oil bath and concentrated by rotary evaporation to yield a brown oil. The reaction vial was fitted with an aluminum–PTFE crimp cap and the crude farnesyl phosphate was further concentrated on hi-vac to remove traces of pyridine and trichloroacetonitrile. The resulting brown gum was dissolved in equal parts anhydrous methanol (550 µl, 1.0 M) and 7 N ammonia solution in methanol (550 µl, 1.0 M). The resulting solution was heated to 55 °C by transferring the reaction vial to an oil bath.

After stirring at this temperature for 12 hours, the reaction vial was removed from the oil bath and concentrated by rotary evaporation to yield a brown gum. This crude product was purified directly by flash column chromatography on silica gel ($CH_2Cl_2$/MeOH = 0/1 to 1/5). Filtration of the dissolved product through a 0.2 micron PTFE syringe filter afforded **1** as a pale yellow oil. This oil was taken up in 1 ml of Milli-Q water, flash frozen and subjected to lyophilization to yield **1** (31 mg, 12% over 3 steps) as a pale yellow solid.

**$^1$H NMR (300 MHz, $CD_3OD$)** δ 5.60 (t, $J$ = 4.4 Hz, 1H), 5.41 (td, $J$ = 6.9, 1.6 Hz, 1H), 5.16 – 5.08 (m, 2H), 4.49 – 4.35 (m, 3H), 3.84 (ddd, $J$ = 7.4, 5.2, 2.8 Hz, 1H), 3.77 (dd, $J$ = 12.4, 2.8 Hz, 1H), 3.69 – 3.59 (m, 2H), 2.17 – 1.97 (m, 8H), 1.73 (q, $J$ = 1.2 Hz, 3H), 1.67 (d, $J$ = 1.4 Hz, 6H), 1.61 (d, $J$ = 1.4 Hz, 3H).

**$^{13}$C NMR (126 MHz, $CD_3OD$)** δ 141.98, 137.87, 133.62, 126.97, 126.63, 124.52 (d), 117.57, 100.27 (d), 86.77, 73.57, 69.95 (d), 64.67 (d), 34.56, 34.15, 28.92, 28.84, 27.20, 24.97, 24.92, 19.00.

**IR** ($cm^{-1}$): 3334, 2963, 2910, 2850, 2107, 1667, 1451, 1372, 1216, 1080, 1011

**HRMS (ESI):** calc. for $C_{20}H_{33}N_3O_7P^-$ ($M^-$) 458.2061, found 458.2060.

**3-azido-3-deoxy-1-*O*-methyl-3,5-di-*O*-*t*-butyldiphenylsilyl-D-arabinofuranose (SI-5):**

A flame-dried, 250 mL round-bottomed flask equipped with a magnetic stir bar was charged with **SI-4** (5.00 g, 26.43 mmol, 1.0 equiv), sealed with a rubber septum, evacuated and backfilled with nitrogen three times, and placed under an argon atmosphere.[3] **SI-4** was dissolved in anhydrous DMF (88 mL, 0.3 M) and imidazole (7.19 g, 105.72 mmol, 4.0 equiv) was added to the resulting solution as a solid in a single portion. TBDPSCl (15.3 ml, 55.50 mmol, 2.1 equiv) was added to the reaction mixture neat, in a dropwise fashion over the course of 2 minutes. The resulting solution was stirred at room temperature for 6 hours.

After this time, the reaction was quenched by pouring into sat. aq. $NaHCO_3$ (300 mL). The layers were separated and the aqueous layer was extracted with pentane (3 × 300 mL). The combined organic layers were washed with brine (500 mL), dried over anhydrous $Na_2SO_4$, filtered, and concentrated under reduced pressure by rotary evaporation to provide a crude yellow oil. Purification by flash column chromatography on silica gel ($Et_2O$/hexanes = 0/10 to 1/5) afforded **SI-5** (12.90 g, 77%) as a colorless oil.

**$^1$H NMR (400 MHz, $CDCl_3$)** δ 7.73 (ddd, *J* = 8.0, 4.6, 1.6 Hz, 4H), 7.70 – 7.64 (m, 4H), 7.53 – 7.38 (m, 12H), 4.75 (d, *J* = 1.1 Hz, 1H), 4.22 (dd, *J* = 2.8, 1.1 Hz, 1H), 4.02 (td, *J* = 5.8, 4.7 Hz, 1H), 3.95 – 3.81 (m, 3H), 3.17 (s, 3H), 1.13 (s, 9H), 1.10 (s, 9H)

**$^{13}$C NMR (101 MHz, $CDCl_3$)** δ 135.78, 135.66, 135.62, 133.24, 133.22, 132.93, 132.65, 130.11, 130.03, 129.82, 129.79, 127.92, 127.83, 127.80, 127.76, 109.05, 82.29, 82.25, 68.42, 64.01, 54.90, 26.87, 26.85, 19.34, 19.10

**IR** ($cm^{-1}$): 3072, 3046, 2953, 2930, 2893, 2860, 2101, 1587, 1468, 1425, 1385, 1359, 1263, 1110

**HRMS (ESI):** calc. for $C_{38}H_{51}N_4O_4Si_2^+$ ($M + NH_4^+$): 683.3443, found 683.3441

**1-*O*-acetyl-3-azido-3-deoxy-3,5-di-*O*-*t*-butyldiphenylsilyl-D-arabinofuranose (SI-6).**

A flame-dried, 250 mL round-bottomed flask equipped with a magnetic stir bar was charged with **SI-5** (5.00 g, 7.32 mmol, 1.0 equiv), sealed with a rubber septum, evacuated and backfilled with nitrogen three times, and placed under an argon atmosphere. **SI-5** was dissolved in $Ac_2O$ (15 mL, 0.5 M). The resulting solution was cooled to 0 °C by transferring the reaction apparatus to an ice-water bath. After stirring for 5 minutes, a solution of $H_2SO_4$ in $Ac_2O$ (200 µl in 3.0 ml) was added to the reaction mixture in a dropwise fashion over the course of 2 minutes. The reaction apparatus was removed from the ice-water bath and was allowed to warm to room temperature.

After stirring for 45 minutes, the reaction was quenched through the addition of solid $Na_2CO_3$. Once bubbling of the mixture ceased upon $Na_2CO_3$ addition, the reaction was poured into sat. aq. $NaHCO_3$ (300 mL). The layers were separated and the aqueous layer was extracted with $Et_2O$ (3 × 300 mL). The combined organic layers were washed with brine (500 mL), dried over anhydrous $Na_2SO_4$, filtered, and concentrated under reduced pressure by rotary evaporation to provide a crude colorless oil. Purification by flash column chromatography on silica gel ($Et_2O$/hexanes = 0/10 to 1/5) afforded **SI-6** (4.51 g, 89%. α:β = 1.0:1.7) as a colorless oil.

**$^1$H NMR (300 MHz, $CDCl_3$)** δ 7.78 – 7.58 (m, 8H), 7.53 – 7.32 (m, 12H), 6.03 (d, *J* = 1.1 Hz, 0.62H, H-1α), 5.55 (d, *J* = 3.6 Hz, 0.38H, H-1β), 4.23 (ddd, *J* = 9.2, 3.6, 1.8 Hz, 1.3H), 4.09 (dt, *J* = 6.3, 4.7 Hz, 0.61H), 4.00 (dd, *J* = 5.2, 2.8 Hz, 0.61H), 3.94 – 3.79 (m, 1.22H), 3.79 – 3.68 (m, 1.01H), 1.89 (s, 1.60H), 1.86 (s, 1.21H), 1.11 (s, 3H), 1.09 (s, 4.3H), 1.07 (s, 6.1H).

**IR** ($cm^{-1}$): 3072, 3042, 2956, 2926, 2890, 2857, 2104, 1750, 1587, 1471, 1428, 1355, 1223, 1134, 1007

**1-*O*-diallylphosphoryl-3-azido-3-deoxy-3,5-di-*O*-*t*-butyldiphenylsilyl-D-arabinofuranose (SI-7).**

A flame-dried, 10 mL round-bottomed flask equipped with a magnetic stir bar was charged with **SI-6** (1.00 g, 1.44 mmol, 1.0 equiv), sealed with a rubber septum, evacuated and backfilled with nitrogen three times, and placed under an argon atmosphere. **SI-6** was dissolved in anhydrous $CH_2Cl_2$ (2.9 ml, 0.5 M) and the resulting solution was cooled to 0 °C by transferring the reaction apparatus to an ice-water bath. After stirring at this temperature for 5 min, 33% HBr in AcOH (529 µl, 4.10 mmol, 2.0 equiv) was added to the reaction flask in a dropwise fashion. Following this addition, the reaction apparatus was immediately removed from the ice-water bath and allowed to equilibrate to room temperature.

After stirring for 1 hour, the stir bar was removed from the reaction flask and the reaction mixture was concentrated *in vacuo*. The resulting orange gum was dried on hi-vac for 1 hour. During this interim period, a second 25 mL round-bottomed flask equipped with a magnetic stir bar and 4 Å molecular sieves (1.00 g) was sealed with a rubber septum and evacuated. The flask was flame dried 3 times to activate the molecular sieves. Upon cooling, the reaction apparatus was evacuated and backfilled with nitrogen three times, and placed under an argon atmosphere. Diallylphosphate (513 mg, 4.10 mmol, 2.0 equiv) was transferred to the reaction flask in a single portion and dissolved in anhydrous toluene (2.9 ml, 0.5 M). Triethylamine (600 µl, 4.32 mmol, 3 equiv) was added to the reaction mixture in a dropwise fashion over the course of 5 minutes, after which the reaction was cooled to -10 °C by transferring the reaction apparatus to a NaCl-ice bath. After stirring at this temperature for 5 minutes, the glycosyl bromide intermediate was dissolved in anhydrous toluene (2.9 ml, 0.5 M). The resulting solution was transferred to the reaction flask in a dropwise fashion. Following this final addition, the reaction flask was allowed to slowly warm to room temperature.

After stirring for 10 hours, the reaction was quenched by pouring into sat. aq. $NH_4Cl$ (150 mL). The layers were separated and the aqueous layer was extracted with $Et_2O$ (3 × 1500 mL). The combined organic layers were washed with brine (300 mL), dried over anhydrous $Na_2SO_4$, filtered, and concentrated under reduced pressure by rotary evaporation to provide a crude oil. Purification by flash column chromatography on silica gel (EtOAc/hexanes = 0/10 to 1/2) afforded **SI-7** (701 mg, 60% over 2 steps) as a colorless oil.

**$^1$H NMR (400 MHz, $C_6D_6$)** δ 7.84 (dt, *J* = 8.1, 1.5 Hz, 5H), 7.81 – 7.70 (m, 6H), 7.34 – 7.18 (m, 14H), 6.26 (d, *J* = 4.8 Hz, 0.33H), 5.88 – 5.55 (m, 3H), 5.13 (ddt, *J* = 32.8, 17.2, 1.6 Hz, 3H), 5.00 – 4.83 (m, 3H), 4.63 (d, *J* = 2.3 Hz, 0.33H), 4.57 – 4.21 (m, 5H), 4.14 – 4.02 (m, 2H), 3.93 – 3.77 (m, 2H), 3.72 (q, *J* = 6.3 Hz, 0.66H), 1.24 (s, 6H), 1.17 (d, *J* = 3.1 Hz, 10H), 1.10 (s, 2H).

**$^{13}$C NMR (101 MHz, $C_6D_6$)** δ 135.98, 135.84, 135.78, 135.70, 135.67, 135.64, 133.14, 133.10, 133.05, 132.88, 132.80, 132.77, 132.73, 132.68, 132.58, 132.18, 130.25, 130.19, 130.16, 129.87, 129.84, 117.29, 117.05, 105.70, 105.64, 98.84, 98.78, 84.51, 80.73, 77.64, 77.56, 68.02, 67.85, 67.80, 67.71, 67.68, 67.53, 67.49, 66.67, 65.87, 63.74, 26.75, 26.71, 26.69, 26.67, 19.19, 19.15, 19.11, 18.89.

**IR** ($cm^{-1}$): 3073, 3046, 2949, 2930, 2890, 2857, 2104, 1591, 1471, 1428, 1269, 1113, 1020, 931, 818

**1-*O*-(*Z,Z*)-farnesylphosphoryl-3-azido-3-deoxy-β-D-arabinofuranose, 3-AzFPA (2).**

A flame-dried, 5 mL microwave vial equipped with a magnetic stir bar and was charged with **SI-7** (300 mg, 0.37 mmol, 1.0 equiv), sealed with a rubber septum, evacuated and backfilled with nitrogen three times, and placed under an argon atmosphere. **SI-7** was dissolved in anhydrous $CH_2Cl_2$ (1.4 mL, 0.4 M) and $Pd(PPh_3)_4$ (4 mg, 1 mol%) was added to the reaction vial in a single portion. Following this addition phenylsilane (90 µl, 0.74 mmol, 2.0 equiv) was added to the reaction vial neat, in a dropwise fashion over the course of 2 minutes.

After stirring for 3 hours, the contents of the reaction vial were concentrated *in vacuo*. The reaction vial was fitted with an aluminum–PTFE crimp cap and dried on hi-vac for 1 hour to remove trace solvent. Following this, the reaction vial was evacuated and backfilled with nitrogen three times, and placed under an argon atmosphere. The crude arabinfuranosyl phosphate was dissolved in anhydrous pyridine (740 µl, 0.5 M). (*Z,Z*)-farnesol (247 mg, 1.10 mmol, 3.0 equiv) and trichloroacetonitrile (370 µl, 3.70 mmol, 10.0 equiv) were added to the reaction vial neat, in single portions. The resulting solution was heated to 40 °C by transferring the reaction vial to an oil bath.

After stirring at this temperature for 12 hours, the reaction vial was removed from the oil bath and concentrated by rotary evaporation to yield a brown oil. The reaction vial was fitted with a rubber septum and the crude farnesyl phosphate was further concentrated on hi-vac to remove traces of pyridine and trichloroacetonitrile. The resulting brown gum was dissolved in equal parts anhydrous methanol (550 µl, 1.0 M) and 7 N ammonia solution in methanol (550 µl, 1.0 M). Ammonium fluoride (137 mg, 3.70 mmol, 10.0 equiv) was added to the reaction vial in a single portion and the vial was fitted with an aluminum–PTFE crimp cap. The resulting solution was heated to 55 °C by transferring the reaction vial to an oil bath.

After stirring at this temperature for 12 hours, the reaction vial was removed from the oil bath and concentrated by rotary evaporation to yield a brown gum. This crude product was purified directly by flash column chromatography on silica gel ($CH_2Cl_2$/MeOH = 0/1 to 1/5). Filtration of the dissolved product through a 0.2 micron PTFE syringe filter afforded **2** as a pale brown oil. This oil was taken up in 1 ml of Milli-Q water, flash frozen and subjected to lyophilization to yield **2** (25 mg, 15% over 3 steps) as a pale yellow solid.

**$^1$H NMR (400 MHz, $CD_3OD$)** δ 5.50 (t, *J* = 4.2 Hz, 1H), 5.44 (t, *J* = 6.9 Hz, 1H), 5.20 – 5.12 (m, 2H), 4.46 (t, *J* = 6.7 Hz, 2H), 4.13 – 4.01 (m, 2H), 3.82 – 3.72 (m, 2H), 3.68 – 3.61 (m, 1H), 2.18 – 2.05 (m, 8H), 1.77 (d, *J* = 1.5 Hz, 3H), 1.71 (s, 7H), 1.65 (d, *J* = 1.3 Hz, 3H).

**$^{13}$C NMR (101 MHz, $CD_3OD$)** δ 139.47, 135.22, 130.99, 124.38, 124.00, 121.93, 121.85, 97.27, 97.21, 81.24, 76.95, 63.95, 62.36, 62.05, 62.00, 31.91, 31.52, 26.27, 26.21, 24.56, 23.66, 22.32, 22.28, 16.35.

**$^{31}$P NMR (162 MHz, $CD_3OD$)** δ -0.74.

**IR** ($cm^{-1}$): 3234, 2963, 2920, 2847, 2091, 1670, 1481, 1372, 1276, 1213, 1054, 1021, 937

**HRMS (ESI):** calc. for $C_{20}H_{33}N_3O_7P^-$ ($M^-$) 458.2061, found 458.2040.

**5-azido-5-deoxy-1-*O*-methyl-2,3-di-*O*-*t*-butyldiphenylsilyl-D-arabinofuranose (SI-9):**

A flame-dried, 250 mL round-bottomed flask equipped with a magnetic stir bar was charged with **SI-8** (5.00 g, 26.43 mmol, 1.0 equiv), sealed with a rubber septum, evacuated and backfilled with nitrogen three times, and placed under an argon atmosphere.[4] **SI-8** was dissolved in anhydrous DMF (88 mL, 0.3 M) and imidazole (7.19 g, 105.72 mmol, 4.0 equiv) was added to the resulting solution as a solid in a single portion. TBDPSCl (15.3 ml, 55.50 mmol, 2.1 equiv) was added to the reaction mixture neat, in a dropwise fashion over the course of 2 minutes. The resulting solution was stirred at room temperature for 12 hours.

After this time, the reaction was quenched by pouring into sat. aq. $NaHCO_3$ (300 mL). The layers were separated and the aqueous layer was extracted with pentane (3 × 300 mL). The combined organic layers were washed with brine (500 mL), dried over anhydrous $Na_2SO_4$, filtered, and concentrated under reduced pressure by rotary evaporation to provide a crude yellow oil. Purification by flash column chromatography on silica gel ($Et_2O$/hexanes = 0/10 to 1/5) afforded **SI-9** (15.17 g, 84%) as a colorless oil.

**$^1$H NMR (600 MHz, $CDCl_3$)** δ 7.65 (dd, *J* = 7.4 Hz, 2H), 7.62 – 7.56 (m, 6H), 7.46 – 7.41 (m, 4H), 7.34 – 7.32 (m, 8H), 4.68 (d, *J* = 1.1 Hz, 1H), 4.34 (d, *J* = 1.2 Hz, 1H), 4.03 – 4.01 (m, 2H), 3.12 (s, 3H), 2.82 (dd, 12.2, 7.12 Hz, 1H), 2.62 (d, 13.4, 3.3 Hz, 1H), 1.00 (s, 9H), 0.99 (s, 9H)

**$^{13}$C NMR (151 MHz, $CDCl_3$)** δ 135.82, 135.79, 135.77, 133.62, 133.30, 132.99, 132.76, 129.98, 129.96, 129.91, 127,89, 127.82, 127.78, 109.45, 85.39, 83.59, 81.25, 54.48, 52,52, 26.86, 26.78, 19.21, 19.12

**IR** (cm$^{-1}$): 3049, 3069, 2959, 2926, 2853, 2098, 1750, 1471, 1428, 1355, 1226, 1007, 954

**1-*O*-acetyl-5-azido-5-deoxy-2,3-di-*O*-*t*-butyldiphenylsilyl-D-arabinofuranose (SI-10).**

A flame-dried, 250 mL round-bottomed flask equipped with a magnetic stir bar was charged with **SI-9** (5.00 g, 7.32 mmol, 1.0 equiv), sealed with a rubber septum, evacuated and backfilled with nitrogen three times, and placed under an argon atmosphere. **SI-9** was dissolved in $Ac_2O$ (15 mL, 0.5 M). The resulting solution was cooled to 0 °C by transferring the reaction apparatus to an ice-water bath. After stirring for 5 minutes, a solution of $H_2SO_4$ in $Ac_2O$ (200 µl in 3.0 ml) was added to the reaction mixture in a dropwise fashion over the course of 2 minutes. The reaction apparatus was removed from the ice-water bath and was allowed to warm to room temperature.

After stirring for 45 minutes, the reaction was quenched through the addition of solid $Na_2CO_3$. Once bubbling of the mixture ceased upon $Na_2CO_3$ addition, the reaction was poured into sat. aq. $NaHCO_3$ (300 mL). The layers were separated and the aqueous layer was extracted with $Et_2O$ (3 × 300 mL). The combined organic layers were washed with brine (500 mL), dried over anhydrous $Na_2SO_4$, filtered, and concentrated under reduced pressure by rotary evaporation to provide a crude colorless oil. Purification by flash column chromatography on silica gel ($Et_2O$/hexanes = 0/10 to 1/5) afforded **SI-10** (3.85 g, 76%) as a colorless oil.

**$^1$H NMR (600 MHz, $CDCl_3$)** δ 7.57 (t, *J* = 6.5 Hz, 4H), 7.51 (d, *J* = 7.4 Hz, 2H), 7.48 (d, *J* = 7.51 Hz, 2H), 7.46 – 7.39 (m, 4H), 7.36 – 7.34 (m, 4H), 7.30 (dt, *J* = 7.5, 3.3 Hz, 4H), 6.06 (s, 1H), 4.29 (s, 1H), 4.18 (dt, *J* = 6.8, 1.9 Hz, 1H), 4.00 (d, *J* = 0.9 Hz, 1H), 3.05 (dd, *J* = 12.4, 7.5 Hz, 1H), 2.71 (dd, *J* = 12.9, 5.0 Hz, 1H), 1.98 (s, 3H), 0.98 (s, 9H), 0.96 (s, 9H).

**1-*O*-diallylphosphoryl-5-azido-5-deoxy-2,3-di-*O*-*t*-butyldiphenylsilyl-D-arabinofuranose (SI-11).**

A flame-dried, 10 mL round-bottomed flask equipped with a magnetic stir bar was charged with **SI-10** (1.00 g, 1.44 mmol, 1.0 equiv), sealed with a rubber septum, evacuated and backfilled with nitrogen three times, and placed under an argon atmosphere. **SI-10** was dissolved in anhydrous $CH_2Cl_2$ (2.9 ml, 0.5 M) and the resulting solution was cooled to 0 °C by transferring the reaction apparatus to an ice-water bath. After stirring at this temperature for 5 min, 33% HBr in AcOH (529 μl, 4.10 mmol, 2.0 equiv) was added to the reaction flask in a dropwise fashion. Following this addition, the reaction apparatus was immediately removed from the ice-water bath and allowed to equilibrate to room temperature.

After stirring for 1 hour, the stir bar was removed from the reaction flask and the reaction mixture was concentrated *in vacuo*. The resulting orange gum was dried on hi-vac for 1 hour. During this interim period, a second 25 mL round-bottomed flask equipped with a magnetic stir bar and 4 Å molecular sieves (1.00 g) was sealed with a rubber septum and evacuated. The flask was flame dried 3 times to activate the molecular sieves. Upon cooling, the reaction apparatus was evacuated and backfilled with nitrogen three times, and placed under an argon atmosphere. Diallylphosphate (513 mg, 4.10 mmol, 2.0 equiv) was transferred to the reaction flask in a single portion and dissolved in anhydrous toluene (2.9 ml, 0.5 M). Triethylamine (600 μl, 4.32 mmol, 3 equiv) was added to the reaction mixture in a dropwise fashion over the course of 5 minutes, after which the reaction was cooled to -10 °C by transferring the reaction apparatus to a NaCl-ice bath. After stirring at this temperature for 5 minutes, the glycosyl bromide intermediate was dissolved in anhydrous toluene (2.9 ml, 0.5 M). The resulting solution was transferred to the reaction flask in a dropwise fashion. Following this final addition, the reaction flask was allowed to slowly warm to room temperature.

After stirring for 10 hours, the reaction was quenched by pouring into sat. aq. $NH_4Cl$ (150 mL). The layers were separated and the aqueous layer was extracted with $Et_2O$ (3 × 1500 mL). The combined organic layers were washed with brine (300 mL), dried over anhydrous $Na_2SO_4$, filtered, and concentrated under reduced pressure by rotary evaporation to provide a crude oil. Purification by flash column chromatography on silica gel (EtOAc/hexanes = 0/10 to 1/2) afforded **SI-11** (798 mg, 60% over 2 steps) as a colorless oil.

**$^1$H NMR (300 MHz, $CD_3OD$)** δ 7.71 – 7.61 (m, 6H), 7.62 – 7.60 (m, 2H), 7.46 – 7.42 (m, 4H), 7.40 – 7.33 (m, 8H), 5.87 – 5.74 (m, 2H), 5.35 – 5.16 (m, 2H), 4.54 – 4.34 (m, 3H), 3.84 – 3.80 (m, 1H), 3.04 (dd, *J* = 13.6, 10.26 Hz, 1H), 2.08 (dd, *J* = 12.8, 3.7 Hz, 1H), 1.10 (s, 9H), 1.01 (s, 9H).

**IR** ($cm^{-1}$): 3072, 3046, 2959, 2929, 2889, 2853, 2094, 1468, 1425, 1276, 1164, 1100, 1021

**1-*O*-(*Z*,*Z*)-farnesylphosphoryl-5-azido-5-deoxy-β-D-arabinofuranose, 5-AzFPA (3).**

A flame-dried, 5 mL microwave vial equipped with a magnetic stir bar and was charged with **SI-11** (300 mg, 0.37 mmol, 1.0 equiv), sealed with a rubber septum, evacuated and backfilled with nitrogen three times, and placed under an argon atmosphere. **SI-11** was dissolved in anhydrous $CH_2Cl_2$ (1.4 mL, 0.4 M) and $Pd(PPh_3)_4$ (4 mg, 1 mol%) was added to the reaction vial in a single portion. Following this addition phenylsilane (90 µl, 0.74 mmol, 2.0 equiv) was added to the reaction vial neat, in a dropwise fashion over the course of 2 minutes.

After stirring for 3 hours, the contents of the reaction vial were concentrated *in vacuo*. The reaction vial was fitted with an aluminum–PTFE crimp cap and dried on hi-vac for 1 hour to remove trace solvent. Following this, the reaction vial was evacuated and backfilled with nitrogen three times, and placed under an argon atmosphere. The crude arabinfuranosyl phosphate was dissolved in anhydrous pyridine (740 µl, 0.5 M). (*Z*,*Z*)-farnesol (247 mg, 1.10 mmol, 3.0 equiv) and trichloroacetonitrile (370 µl, 3.70 mmol, 10.0 equiv) were added to the reaction vial neat, in single portions. The resulting solution was heated to 40 °C by transferring the reaction vial to an oil bath.

After stirring at this temperature for 12 hours, the reaction vial was removed from the oil bath and concentrated by rotary evaporation to yield a brown oil. The reaction vial was fitted with a rubber septum and the crude farnesyl phosphate was further concentrated on hi-vac to remove traces of pyridine and trichloroacetonitrile. The resulting brown gum was dissolved in equal parts anhydrous methanol (550 µl, 1.0 M) and 7 N ammonia solution in methanol (550 µl, 1.0 M). Ammonium fluoride (137 mg, 3.70 mmol, 10.0 equiv) was added to the reaction vial in a single portion and the vial was fitted with an aluminum–PTFE crimp cap. The resulting solution was heated to 55 °C by transferring the reaction vial to an oil bath.

After stirring at this temperature for 12 hours, the reaction vial was removed from the oil bath and concentrated by rotary evaporation to yield a brown gum. This crude product was purified directly by flash column chromatography on silica gel ($CH_2Cl_2$/MeOH = 0/1 to 1/5). Filtration of the dissolved product through a 0.2 micron PTFE syringe filter afforded **3** as a pale brown oil. This oil was taken up in 1 ml of Milli-Q water, flash frozen and subjected to lyophilization to yield **3** (31 mg, 18% over 3 steps) as a pale yellow solid.

**$^1$H NMR (600 MHz, $CD_3OD$)** δ 5.55 (dd, $J$ = 6.6, 3.3 Hz, 1H), 5.42 (t, $J$ = 5.9 Hz, 1H), 5.16 – 5.10 (m, 2H), 4.45 (t, $J$ = 7.0 Hz, 2H), 4.00 – 3.95 (m, 2H), 3.86 – 3.81 (m, 2H), 3.54 – 3.49 (m, 1H), 2.16 – 1.96 (m, 8H), 1.77 (s, 3H), 1.67 (s, 6H), 1.61 (s, 3H).

**IR** ($cm^{-1}$): 3234, 2963, 2920, 2847, 2091, 1670, 1451, 1372, 1276, 1213, 1137, 1154, 1054, 1021

**HRMS (ESI):** calc. for $C_{20}H_{33}N_3O_7P^-$ ($M^-$) 458.2061, found 458.2037.
